# A Scalable Framework for Species-Resolved Human Gut Microbiome Profiling Using Full-Length 16S rRNA Sequencing

**DOI:** 10.64898/2026.06.21.732309

**Authors:** Parth Sarin, Paras Sehgal, Vasudev Paveri, Sakshi Rai, Ashriti Chettri, Rahul C. Bhoyar, Shahzad Mirza, Rajesh Karkaryate, Sourav Sen Gupta, JP Devraj, Sridhar Sivasubbu

## Abstract

The gut microbiota plays a fundamental role in human health, nutrition, immune development, and disease, driving widespread adoption of 16S rRNA gene sequencing for microbial community characterization. Short-read V3–V4 sequencing remains the dominant approach for large-scale microbiome studies; however, interrogation of only a small fraction of the 16S gene limits phylogenetic resolution and frequently restricts biological interpretation at the species level. Although full-length (V1–V9) 16S sequencing has emerged as a promising alternative, comprehensive evaluation of highly multiplexed full-length workflows in complex human gut microbiomes remains limited. Here, we establish and evaluate a full-length 16S framework for species-resolved human gut microbiome profiling. The workflow was assessed using defined microbial communities, technical replicates, and healthy human fecal microbiomes. Full-length sequencing generated highly concordant taxonomic profiles across independent technical workflows and enabled reproducible recovery of complex microbial communities at both genus and species levels. Application to human fecal microbiomes revealed substantial inter-individual heterogeneity together with extensive ASV-level microdiversity, highlighting the ability of full-length sequencing to resolve fine-scale phylogenetic variation within dominant gut-associated taxa. To quantify the analytical gain afforded by full-length sequencing, V3–V4 datasets were computationally reconstructed directly from identical full-length reads, eliminating methodological and biological confounders. While alpha diversity metrics and overall community structure remained highly concordant between approaches, full-length sequencing markedly improved taxonomic resolution, increasing species-level assignment from approximately 20% to 98% and resolving substantial intra-genus diversity within clinically and ecologically relevant genera including Bifidobacterium, Prevotella, Blautia, Enterococcus, and Klebsiella. Collectively, these findings position full-length 16S sequencing as an enabling technology for the next generation of microbiome studies, where species-level resolution can be integrated with large-scale cohort, longitudinal, and population-health investigations.

## 1. Introduction

Microbial communities influence host physiology, ecological dynamics, and disease processes, needing precise, scalable, and high-resolution taxonomic classification (1–3). The 16S rRNA gene (∼1,500 bp) is a key marker for bacterial phylogeny and taxonomy (4–6). It consists of conserved sections and nine hypervariable domains (V1-V9)(7–9). Conserved regions allow for the development of generally applicable primers, whereas hypervariable regions accrue sequence diversity, making it easier to distinguish across taxa(10–12). This dual structural arrangement underpins its long-term value in microbial ecology and clinical microbiology(6,13,14).

The evolution of sequencing technologies has profoundly shaped 16S rRNA-based microbiome studies (15,16) . Early high-throughput efforts using pyrosequencing platforms developed by Roche and semiconductor-based sequencing from Thermo Fisher Scientific enabled initial large-scale microbial surveys, albeit with limitations in read length and accuracy(17–19). The subsequent dominance of Illumina short-read sequencing established a cost-effective and highly scalable framework for microbiota profiling, primarily targeting subregions such as V3–V4 or V4 (16,20,21). While these approaches provide reliable genus-level resolution, numerous studies have demonstrated that taxonomic assignments and inferred community structure are strongly dependent on the chosen variable region (11,22,23). Critically, short-read sequencing often fails to resolve closely related species, resulting in collapsed taxonomic classifications and reduced ecological and clinical interpretability (6,22,24).

Recent advances in third-generation sequencing technologies have enabled full-length 16S rRNA profiling, capturing all variable regions within a single contiguous read(25,26). Platforms such as Oxford Nanopore Technologies(ONT) offer long-read capability and real time sequencing, but historically have been limited by higher raw error rates(27,28). In contrast, Pacific Biosciences HiFi sequencing, based on circular consensus generation, produces highly accurate long reads that substantially improve taxonomic resolution and enable robust species-level classification(6,26,29). This enhanced resolution allows more precise characterization of complex microbial communities and strengthens downstream ecological and association analyses (6,26,30).

To further address limitations in throughput and cost, the PacBio Kinnex 16S workflow incorporates a MASl1lSeql1lbased concatenation strategy, in which multiple fulll1llength amplicons are linked and sequenced as a single molecule, followed by computational deconvolution (31–33). This approach increases effective sequencing yield and supports highly multiplexed study designs while maintaining the high accuracy of HiFi reads (31,33). Although Kinnex and related MASl1lSeq chemistries have so far been primarily demonstrated in relatively lowl1lbiomass or lowl1lcomplexity systems, including fruitl1lfly microbiome work from the Mason laboratory and others (34,35) their performance in high-biomass samples has not been systematically evaluated, particularly in the context of the human gut microbiota.

Collectively, these advances extend 16Sl1lbased microbiota profiling from genusl1llevel surveys toward routine, high resolution species level characterization, particularly in complex and highl1lbiomass ecosystems (6,26,36,37). This study represents the first application of Kinnex-enabled full-length 16S rRNA sequencing in high-biomass samples, establishing a scalable and high resolution framework for microbiota profiling with respect to human gut.

## 2. Materials and Methods

### 2.1 Sample Collection and Cohort

A total of 15 fecal samples were collected from apparently healthy children aged 18–59 months attending community centers in suburban regions of Hyderabad, representing a low- to middle-income population. Samples were obtained in sterile 50 mL containers (#510030, Tarsons, India), transported on dry ice, and stored at −80 °C until further analysis. The study was approved by the Institutional Ethics Committee of ICMR–National Institute of Nutrition(IEC Approval No.CR/4/III/2024)

### 2.2 DNA Extraction and Quality Control

DNA extractions were performed using the QIAamp PowerFecal Pro DNA Kit (#51804, Qiagen, Hilden, Germany), as per manufacturer’s protocol with bead beating for lysis. QC of the eluted DNA was assessed on the Eppendorf Biospectrophotometer using μcuvette 1.0.

### 2.3 16S rRNA Amplification and Kinnex formation

Full-length 16S rRNA genes (V1–V9) were amplified from high-quality genomic DNA using uniquely barcoded primers, enabling robust sample-level indexing and accurate downstream demultiplexing. PCR amplification was performed under optimized conditions to ensure uniform amplification across diverse microbial templates while minimizing bias. The resulting barcoded amplicons from individual samples were quantified and pooled in equimolar proportions. These pooled amplicons were subsequently subjected to a controlled concatenation process to generate long Kinnex arrays (∼18–19 kb), thereby improving sequencing efficiency and throughput on long-read platforms.

### 2.4 Library Preparation and PacBio Sequencing

The concatenated amplicon arrays underwent enzymatic processing, including DNA damage repair and nuclease treatment, to enhance molecular integrity and remove potential artifacts. The processed DNA was then converted into SMRTbell libraries via adapter ligation, making them compatible with single-molecule real-time (SMRT) sequencing. Bead-based purification and cleanup steps were performed to eliminate excess reagents and to enrich appropriately sized library fragments.

The workflow supports high-throughput multiplexing, enabling the simultaneous sequencing of hundreds to over a thousand uniquely barcoded samples within a single run. Final libraries were sequenced on PacBio Revio, leveraging long-read sequencing to generate highly accurate, full-length 16S rRNA sequences. This approach enables high-resolution taxonomic profiling and comprehensive characterization of complex microbial communities.

### 2.5 Bioinformatics Pipeline

Raw sequencing data generated using PacBio single-molecule real-time (SMRT) sequencing were first processed to generate high-accuracy circular consensus sequencing (CCS/HiFi) reads, leveraging multiple passes of individual DNA molecules to achieve high per-base accuracy and substantially reduce random sequencing errors. Stringent quality thresholds were applied during CCS generation to retain only high-confidence reads suitable for downstream analysis. Given the concatenated architecture of Kinnex libraries, the initial processing step involved segmentation of long HiFi reads using skera, which accurately splits concatenated molecules into individual full-length 16S rRNA amplicon units. This segmentation is essential to deconvolute synthetic concatemer structures and ensure that each resulting read corresponds to a single biological amplicon, thereby preserving the integrity of downstream analyses.

Following segmentation, demultiplexing was performed using lima, which identifies and validates barcode sequences present at both ends of each amplicon. This step enables precise assignment of reads to their respective samples while simultaneously trimming barcode and adapter sequences. Additionally, reads with incomplete, asymmetric, or low-confidence barcode matches were filtered out, ensuring high specificity in sample assignment and minimizing cross-sample contamination. Primer sequences were subsequently detected and trimmed, and only reads containing the expected full-length 16S regions were retained, further enhancing dataset quality and consistency.

The processed and curated reads were then analyzed using the HiFi-16S-workflow implemented in Nextflow (https://github.com/PacificBiosciences/HiFi-16S-workflow), which provides a reproducible, modular, and scalable framework for downstream analysis. This workflow integrates multiple steps including stringent quality filtering, error correction, and denoising to resolve amplicon sequence variants (ASVs) at single-nucleotide resolution, thereby overcoming the limitations of traditional OTU-based clustering approaches. The pipeline also incorporates chimera detection and removal to eliminate PCR-derived artifacts, improving the biological accuracy of the dataset. The resulting high-confidence ASVs provide a robust foundation for downstream taxonomic classification, phylogenetic analysis, and comprehensive profiling of microbial community structure and diversity. The resulting high-quality ASVs were used for downstream taxonomic assignment and microbial diversity analyses. This integrated workflow, combining SMRT sequencing with standardized processing tools, enables accurate and high-resolution characterization of complex microbial communities from full-length 16S rRNA gene data.

### 2.6 Validation and Proof-of-Concept Assessment

To assess the performance of the experimental and computational workflow, validation was carried out using defined positive-control microbial communities composed of well-characterized reference strains, including *Escherichia coli (ATCC:25922)* (A), *Klebsiella pneumoniae (ATCC:700603)* (B), *Proteus mirabilis (ATCC:25933)* (C), *Pseudomonas* spp. (ATCC:27853) (D), *Staphylococcus epidermidis (ATCC:35384)* (E), *Enterococcus faecalis* (ATCC: 29212) (F), and *Staphylococcus aureus* (ATCC:25923) (G). A total of 18 experimental conditions were designed to evaluate performance across varying levels of community complexity. These included individual strains (A–G), simple combinations (e.g., A+B, C+D, E+F+G), and complex mixtures comprising four (A+B+C+D) and seven (A+B+C+D+E+F+G) organisms. In addition, dilution series experiments were performed to assess sensitivity and dynamic range, wherein dominant taxa were combined with background organisms at defined ratios of 1:10, 1:100, and 1:1000. All control samples were processed using a standardized workflow, including full-length V1–V9 16S rRNA amplification, Kinnex-based library preparation, PacBio SMRT sequencing, and downstream ASV-based analysis. Observed taxonomic profiles were compared with expected compositions to evaluate accuracy in terms of presence/absence and relative abundance.

Reproducibility was assessed using technical replicates processed independently through the complete workflow, with concordance evaluated using correlation-based metrics (pearson) and compositional similarity measures. In parallel, pipeline benchmarking was performed by evaluating key quality control parameters, including read filtering criteria, minimum read depth thresholds, and ASV inference settings, to ensure consistent and reliable performance across samples.

### 2.7 In Silico Reconstruction of the V3–V4 Hypervariable Region

To directly evaluate the contribution of amplicon length to microbiome inference, full-length V1–V9 HiFi reads were computationally trimmed to the V3–V4 hypervariable region. Trimming was performed using Cutadapt (v5.2) with the canonical V3–V4 primer pair 341F (5′-CCTACGGGNGGCWGCAG-3′) and 806R (5′- GACTACHVGGGTATCTAATCC-3′). Reads containing both primer sequences in the expected orientation were retained, and the intervening V3–V4 region was extracted to generate an in silico V3–V4 dataset derived directly from the original full-length sequencing molecules.

The reconstructed V3–V4 reads were subsequently processed independently using the nf-core/ampliseq pipeline (https://github.com/nf-core/ampliseq). Briefly, primer removal and quality filtering were performed using Cutadapt, ASV inference was conducted using DADA2, and taxonomic classification was performed using the same reference database and analytical framework applied to the full-length dataset. Alpha-diversity, beta-diversity, taxonomic composition, and species-level assignment metrics were then compared between full-length and reconstructed V3–V4 datasets.

### 2.8 s Statistical Analyses

#### 2.8.1 Alpha Diversity Analysis

Alpha diversity analyses were performed at the amplicon sequence variant (ASV) level to compare microbial diversity between full-length V1–V9 and V3–V4 16S rRNA datasets. Diversity was quantified using observed ASV richness, Chao1 richness estimator, Shannon diversity index (H, log2-based), and Simpson diversity index (1 − D). For paired comparisons between sequencing approaches, diversity metrics were calculated from rarefied ASV tables and compared using paired Wilcoxon signed-rank test P-values < 0.05 were considered statistically significant

#### 2.8.2 Beta Diversity Analysis

Beta diversity analyses were performed to evaluate differences in microbial community composition between full-length V1–V9 and V3–V4 16S rRNA datasets. Community dissimilarity was quantified using the Jaccard distance metric, which measures differences in taxonomic composition based on the presence or absence of ASVs. Pairwise distance matrices were visualized using Principal Coordinates Analysis (PCoA) to assess patterns of sample clustering and separation between sequencing approaches. Statistical significance of differences in community composition was evaluated using permutational multivariate analysis of variance (PERMANOVA) with 999 permutations. Differences were considered statistically significant at an adjusted P-value < 0.05 using Benjamini–Hochberg false discovery rate (FDR) procedure.

#### 2.8.3 Rarefaction and Sequencing Depth Assessment

To evaluate the influence of sequencing depth on diversity estimates and species detection, rarefaction analyses were performed using the deepest available subsampled read set from each sample as the source read pool. Reads were randomly subsampled without replacement across 50 evenly spaced sequencing depths spanning a predefined seed depth to the maximum read count.

For each sample and sequencing depth, 20 independent rarefaction replicates were generated and averaged to minimize stochastic variation associated with subsampling. Alpha diversity metrics, including observed ASVs, and Shannon diversity, were calculated at each depth. To assess differences among predefined read-depth groups (10,000, 20,000, 25,000, 50,000, 75,000, and 100,000 reads), pairwise two-sided paired Wilcoxon signed-rank tests were performed using samples shared across all depth comparisons.

#### 2.8.4 Species-Level Resolution Assessment

Species-level classification performance was evaluated by comparing taxonomic assignments obtained from full-length V1–V9 and V3–V4 datasets. The proportion of classified features at each taxonomic rank (phylum, class, order, family, genus, and species) was calculated relative to the total number of assigned features. Comparative analyses focused on species-level recovery, taxonomic concordance, and the ability of each sequencing approach to resolve closely related bacterial taxa. Relative abundance profiles of representative genera were further examined to assess agreement and taxonomic resolution between regions.

#### 2.8.5 Rare Taxa Detection Analysis

To assess the influence of sequencing depth on the recovery of low-abundance community members, ASVs were categorized according to their relative abundance within each sample. Rare ASVs were defined as features with relative abundances between 0.01% and 0.1%, whereas abundant ASVs were defined as features with relative abundances ≥ 0.1%(38,39). Total ASVs included all detected features supported by at least one sequencing read. For each rarefaction depth group (10,000, 20,000, 25,000, 50,000, 75,000, and 100,000 reads), the numbers of total, abundant, and rare ASVs were calculated on a per-sample basis. Species-level relative abundances were subsequently averaged across samples within each depth group to evaluate the effect of sequencing depth on taxonomic recovery.

The 25 most abundant species across all depth groups were visualized individually, while the remaining abundant taxa were aggregated into a single “Other abundant” category. The cumulative relative abundance of the rare biosphere was calculated for each depth group and displayed alongside abundant taxa to assess the contribution of low-abundance organisms across sequencing depths. Distributions of ASV counts were visualized using violin plots with overlaid box-and-whisker plots generated using a Gaussian kernel density estimator (bandwidth = 0.35). Mean species-level relative abundance profiles were visualized using stacked bar charts.

Differences in ASV recovery between sequencing-depth groups were assessed using pairwise two-sided paired Wilcoxon signed-rank tests, leveraging the repeated measurements obtained from the same samples across all rarefaction depths. Resulting P-values were adjusted for multiple testing using the Benjamini–Hochberg false discovery rate (FDR) correction across all pairwise comparisons.

## 3. Results

### 3.1 Analytical validation demonstrates accurate taxonomic recovery across defined microbial communities

The analytical performance of the workflow was systematically evaluated using defined mock microbial communities composed of well-characterized reference strains, including *Escherichia coli*, *Klebsiella pneumoniae*, *Proteus mirabilis*, *Pseudomonas* spp., *Staphylococcus epidermidis*, *Enterococcus faecalis*, and *Staphylococcus aureus*. These controls were designed to rigorously assess taxonomic resolution, quantitative accuracy, and detection sensitivity across gradients of community complexity and abundance. Across all experimental conditions, the pipeline demonstrated high concordance between expected and observed community compositions. In single-species controls, each organism was correctly identified at the species level with negligible background noise, confirming high specificity and minimal cross-contamination (Fig 1A). Raw data with respect to the read quality and other run metrics have been attached in Supplementary table S1. In mixed communities, all constituent taxa were consistently detected, with accurate assignment maintained even among phylogenetically related organisms. Notably, closely related taxa within the *Enterobacteriaceae*, such as *E. coli* and *K. pneumoniae*, and within *Staphylococcaceae*, such as *S. aureus* and *S. epidermidis*, were reliably resolved at the species level, underscoring the discriminatory power of full-length 16S rRNA sequencing combined with ASV-based inference (Fig 1B).

**Figure 1:**
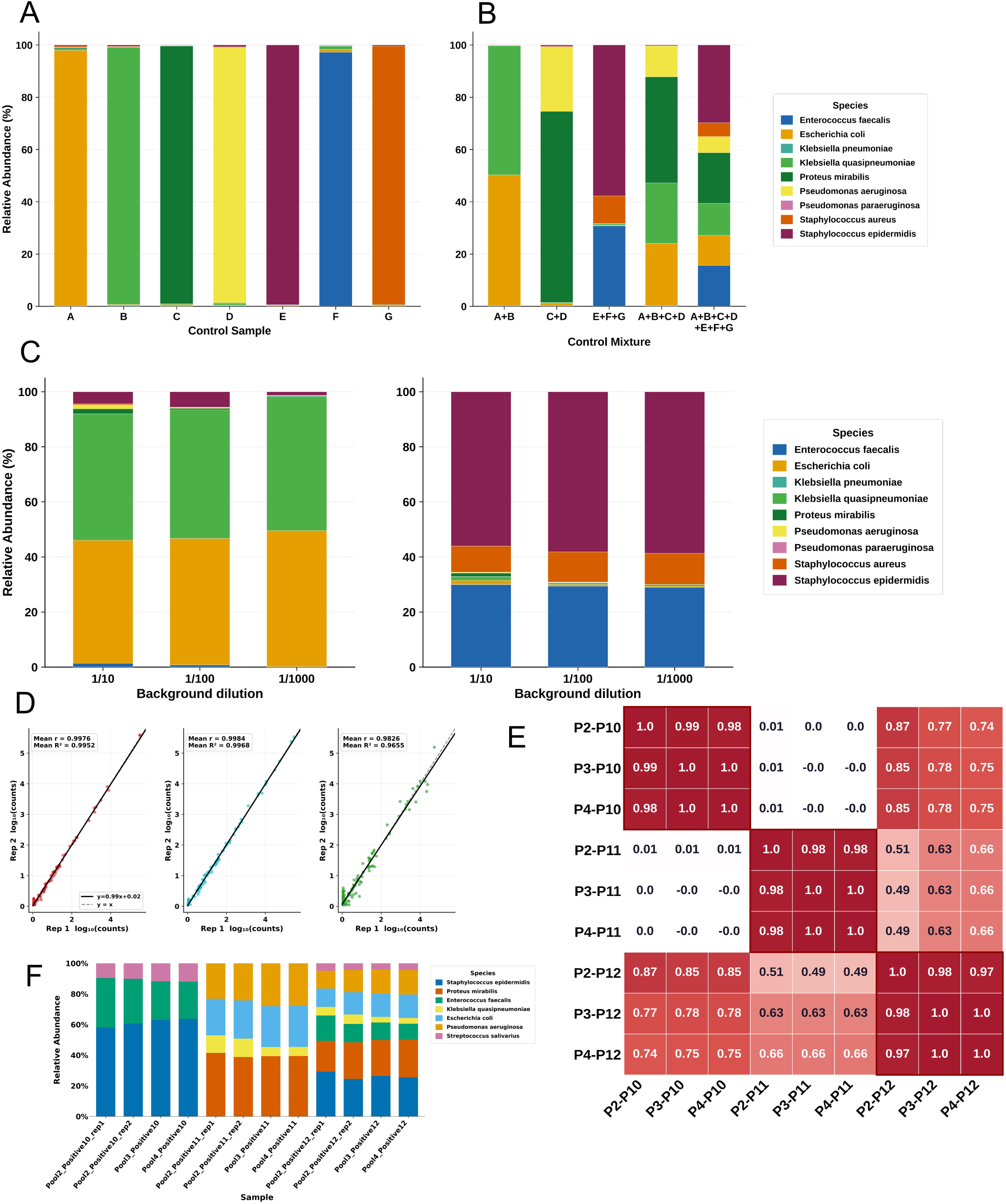
Validation of positive controls using known ATCC strains (observed vs. expected): **(A)** Detection of pure individual isolates for all 7 selected strains; **(B)** Detection of bacterial isolates in mixtures with variable concentrations; **(C)** Assessment of Limit of Detection (LoD) analysis for microbial identification using serial dilutions (1/10, 1/100, and 1/1000); **(D)** Linear regression and Pearson correlation in intra-pool replicability **(E)** Pearson correlation heatmaps showing inter-replicate agreement for positive controls **(F)** Species-level relative abundance profiles of positive controls

Quantitative assessment revealed strong agreement between expected and observed relative abundances across simple and complex mixtures (Fig 1B). Minor deviations were observed, likely attributable to differential amplification efficiencies and inherent PCR biases, but these did not compromise overall compositional fidelity. Dominant taxa such as *E. coli* and *Enterococcus faecalis* in high-abundance mixtures were consistently recovered with proportional representation, while co-occurring taxa were accurately retained without inflation or dropout. Sensitivity analyses using dilution series demonstrated robust detection of low-abundance taxa across a dynamic range spanning three orders of magnitude. Species present at 1/10, 1/100, and 1/1000 relative abundance were reproducibly detected with minimal false positives, indicating high sensitivity and low noise thresholds(Fig 1C). Although stochastic variation increased at extreme dilutions, particularly at the species level, taxonomic assignments remained biologically consistent, supporting the robustness of the pipeline for resolving both dominant and rare members within complex microbial communities.

### 3.2 HiFi Full-length 16SrRNA profiling exhibits high technical reproducibility across library preparations and sequencing runs

Reproducibility was evaluated across two independent axes of technical variation: inter-pool variability arising from library preparation and inter-run variability associated with sequencing. Inter-pool reproducibility was assessed using three samples processed in duplicate across two independent library preparation pools. Pairwise comparisons of species-level relative abundance profiles demonstrated high concordance between pool replicates, with Pearson correlation coefficients approaching unity (Figure 1D), indicating minimal batch effects introduced during library preparation.

Inter-run reproducibility was evaluated using three samples processed in triplicate across independent SMRT sequencing runs. Hierarchical clustering of relative abundance profiles showed tight grouping of replicates by biological identity rather than by sequencing run (Figure 1E), confirming consistency of sequencing performance and downstream processing. Replicate similarity was consistently higher within samples than between samples, indicating preservation of biological signal across runs.

Given that the evaluated samples represent defined equimolar mixtures of reference strains, reproducibility was assessed based on consistency of relative abundance profiles. Across both inter-pool and inter-run comparisons, replicate profiles demonstrated high concordance, with no evidence of systematic variation attributable to library preparation or sequencing (Figures 1D–E). Observed differences in relative abundance were consistent across replicates and did not affect overall compositional recovery. Collectively, these results demonstrate that the workflow yields stable and reproducible community profiles across independent experimental batches.

### 3.3 Application of full-length sequencing to healthy human gut microbiota reveals species-resolved community architecture

To assess performance in complex biological systems, the workflow was applied to a cohort of 15 healthy human fecal samples. Sequencing yielded consistently high read depth per sample, with substantial ASV richness observed across the cohort (Figure 2A), indicating deep sampling of microbial diversity.

**Figure 2.**
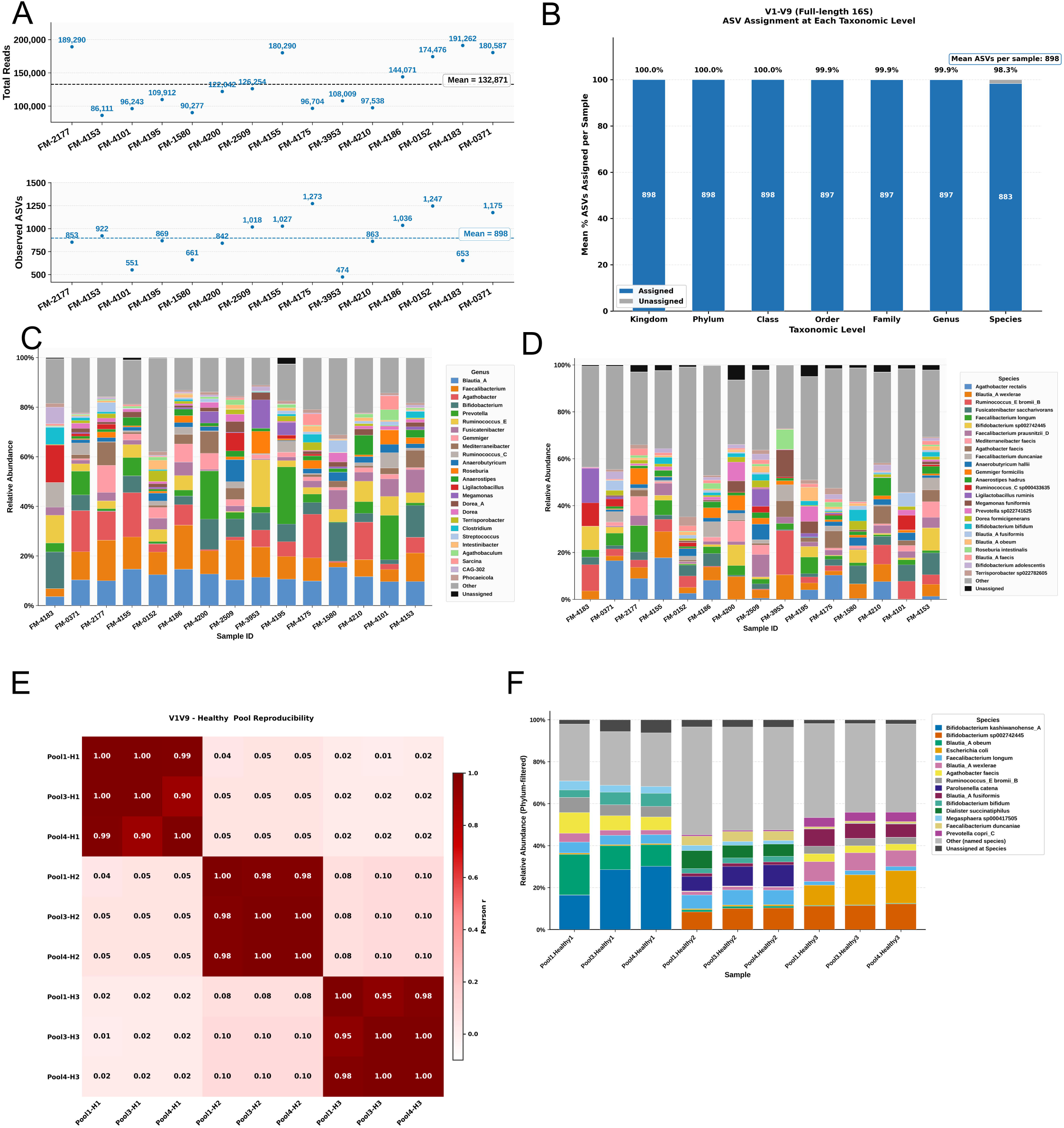
Data from healthy human cohort (real-world samples). **(A)** Total read depth and observed ASV richness per sample generated using Kinnex-based PacBio HiFi CCS sequencing, with mean values indicated. **(B)** Mean and proportions of ASVs assigned and unassigned at every taxonomic level. **(C)** Genus-level relative abundance profiles of 15 healthy samples. **(D)** Species-level relative abundance profiles of 15 healthy samples. **(E)** Pearson correlation heatmaps showing inter-replicate agreement for human samples, respectively. **(F)** Reproducibility species-level relative abundance profiles of human samples in different runs.

Taxonomic assignment rates remained high across hierarchical levels, with the majority of ASVs classified from phylum to species (Figure 2B). As expected, assignment efficiency decreased with increasing taxonomic resolution; however, species-level annotation was retained for a substantial fraction of features, supporting the utility of full-length 16S profiling for high-resolution characterization of the human microbiota.Comprehensive sequencing quality-control metrics, including read filtering and processing statistics, are summarized in Supplementary table S1.

Community composition at the genus level (Figure 2C) recapitulated canonical gut microbiota architecture, with dominance of genera such as Bacteroides, Faecalibacterium, and Prevotella, alongside marked inter-individual variability. Notably, species-level profiling (Figure 2D) revealed extensive intra-genus heterogeneity, with multiple co-occurring species resolved within individual samples. This resolution enabled discrimination of closely related taxa within highly represented genera, overcoming the taxonomic compression inherent to short-read amplicon approaches.

Across samples, species-level profiles exhibited structured diversity rather than stochastic fragmentation, suggesting that the additional resolution captured biologically meaningful variation rather than noise. The ability to resolve co-existing taxa within dominant genera provides a more granular view of community organization, with implications for functional interpretation and downstream comparative analyses.

Collectively, these data demonstrate that full-length 16S sequencing using the described workflow enables high-fidelity reconstruction of gut microbial communities, preserving both macro-scale compositional structure and fine-scale taxonomic diversity within real-world samples.

### 3.4 Reproducibility of microbiota profile in Human Fecal Samples

To further assess reproducibility under biologically complex conditions, three representative human fecal samples were processed across independent sequencing runs. Pairwise comparisons of species-level relative abundance profiles demonstrated near-complete concordance across runs, with Pearson correlation coefficients (R²) approaching unity for all inter-run comparisons (Figure 2E).

Correlation structure was preserved across the full dynamic range of taxa, indicating that both dominant and lower-abundance species were consistently recovered without evidence of run-specific distortion. Replicate profiles retained a highly similar compositional structure, as further illustrated by species-level abundance distributions across runs (Figure 2F), confirming stability of taxonomic assignment and relative abundance estimation in complex microbial communities.

Notably, the high degree of concordance observed in real-world samples mirrors the reproducibility established in defined control experiments, indicating that technical robustness of the workflow is maintained in the presence of substantial biological complexity. These findings establish that the workflow yields highly reproducible microbiota profiles across independent sequencing runs, supporting its application in comparative and longitudinal human cohort studies.

### 3.5 Full-length sequencing preserves ecological structure while expanding species-level resolution relative to V3–V4 profiling

To directly assess the impact of amplicon length on microbiota inference, full-length HiFi reads were computationally truncated to the V3–V4 region using canonical primer coordinates, enabling paired comparisons derived from identical sequencing molecules and thereby eliminating confounding effects of sequencing depth, platform variability, and sample heterogeneity. Across all alpha diversity metrics, full-length (V1–V9) profiles consistently demonstrated significantly higher diversity and richness relative to V3–V4, with marked increases observed in Shannon diversity, Simpson index, and Chao1 estimates (Figure 3A–C; p < 0.001), indicating a systematic underestimation of community complexity by partial-region sequencing. Despite this divergence in diversity, principal component analysis revealed near-complete overlap between V1–V9 and V3–V4 profiles (Figure 3D), with samples clustering by biological identity rather than sequencing region. This concordance establishes that the compositional signals captured by V3–V4 are fully retained within V1–V9, demonstrating that full-length sequencing is non-lossy with respect to overall community structure and that observed differences arise from gain of information rather than loss of signal.

**Figure 3.**
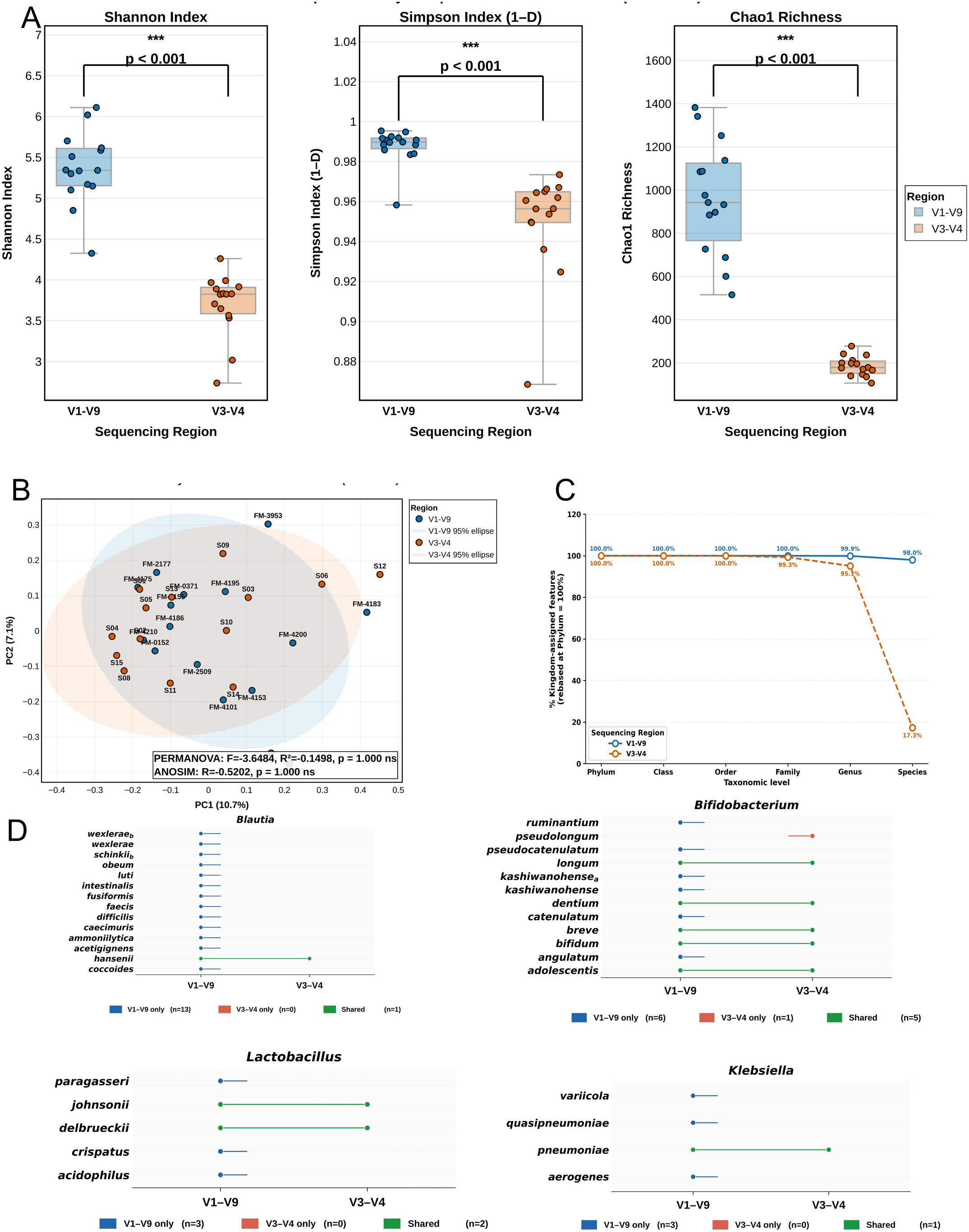
Comparison of microbiome diversity in healthy human real-world samples between the V1-V9 and V3-V4 16S rRNA regions. *(A)* Boxplots showing Shannon diversity, Simpson index, and Chao1 richness for samples sequenced using the full-length V1-V9 region and the V3-V4 region (n = 15 per group). All alpha diversity metrics were significantly higher in V1-V9 compared to V3-V4 (p < 0.001 for all comparisons) (B) Beta diversity evaluated to compare overall community structure between the V1-V9 and V3-V4. (C) Line plot showing the proportion of classified features across taxonomic levels (phylum to species) for V1-V9 and V3-V4 regions of the 16S rRNA gene, normalized to phylum-level assignments (D) Species-level comparison of selected bacterial genera (Prevotella, Blautia, Bifidobacterium, Enterococcus, and Klebsiella) identified using V1-V9 and V3-V4 regions of the 16S rRNA gene.

In contrast, pronounced divergence emerges at finer taxonomic resolution. While taxonomic assignment rates remain comparable at higher hierarchical levels, V3–V4 exhibits a progressive decline in assignability toward lower ranks, culminating in a near-complete loss of species-level resolution (Figure 3E). This reduction manifests as substantial taxonomic compression, wherein multiple distinct species resolved in V1–V9 are collapsed into single or ambiguous assignments in V3–V4 (Figure 3F), particularly within diverse genera such as Prevotella, Blautia, Bifidobacterium, Enterococcus, and Klebsiella. Consequently, intra-genus heterogeneity and species-level diversity are effectively obscured in V3–V4 datasets, limiting biological interpretability and masking functionally and clinically relevant variation. Collectively, these findings demonstrate that while V3–V4 sequencing preserves broad community architecture, it fundamentally truncates diversity and resolution, whereas full-length V1–V9 sequencing captures both global composition and fine-scale taxonomic granularity, providing a substantially more complete and biologically informative representation of microbial communities.

### 3.6. Sequencing Depth Optimization for High Resolution Full-Length Human Gut Microbiota Profiling

To define sequencing depth requirements for full-length V1–V9 microbiome profiling, diversity recovery was evaluated using both rarefaction analysis and explicit read-depth subsampling followed by complete taxonomic reclassification. Rarefaction curves generated from the complete dataset demonstrated rapid accumulation of diversity metrics with progressive saturation at increasing sequencing depths (Figure 4A). Most diversity gains occurred at lower read depths, with observed ASVs approaching approximately 85–90% of their asymptotic values by ∼40K-50K reads/sample, while Shannon diversity reached near-plateau behavior substantially earlier, indicating early stabilization of community evenness and dominant ecological structure.

**Figure 4.**
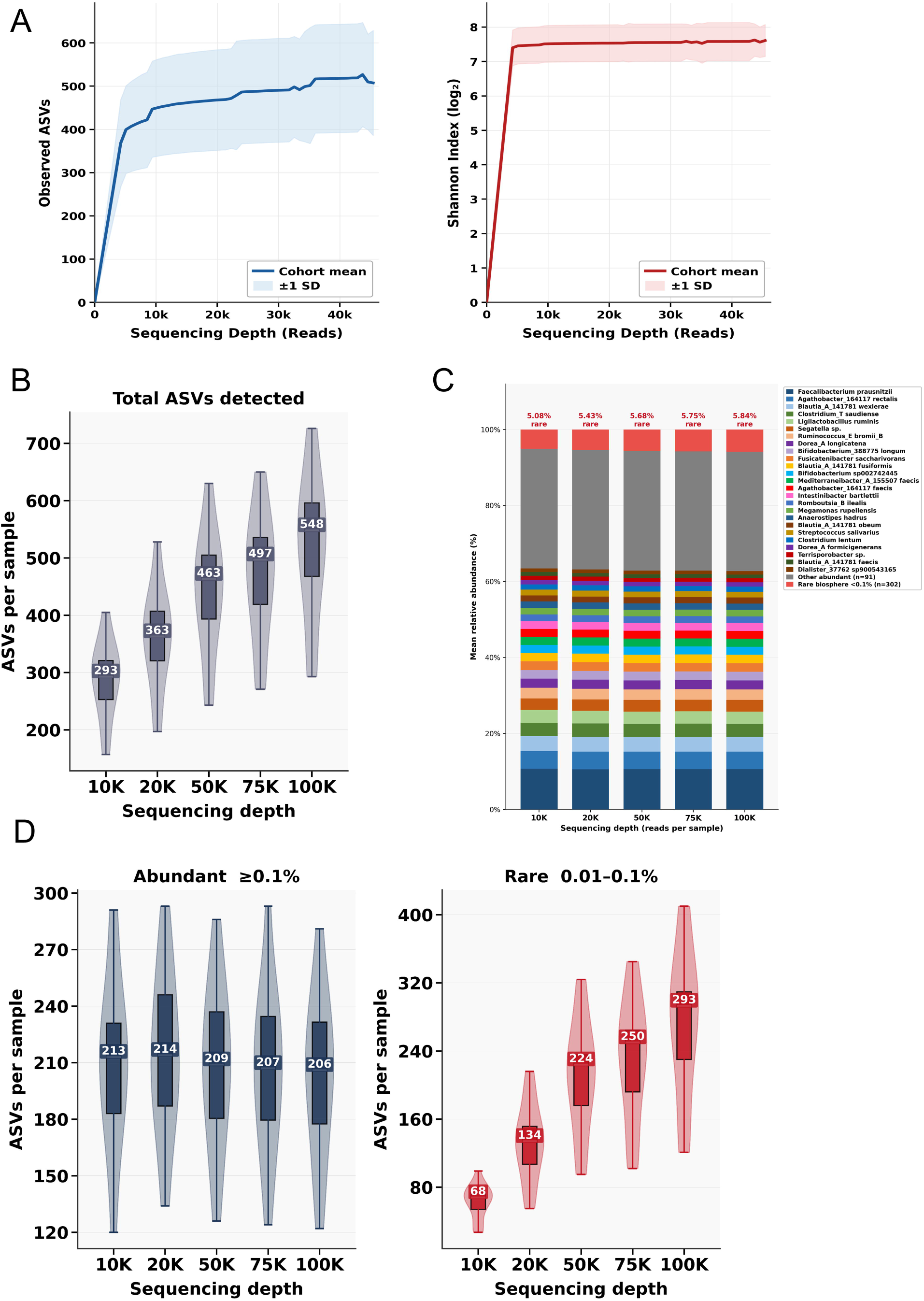
Impact of sequencing depth on diversity and species detection using subsampled read depths. **(A)** Rarefaction curves of alpha diversity metrics (Observed ASVs, Chao1, Shannon, Simpson) across sequencing depths, generated by subsampling the original dataset to defined read depths (10K-100K reads). **(B)** Violin plots with overlaid boxplots showing the distribution of observed ASVs across subsampled read depths (10K-100K reads) using DADA2. **(C)** Relative abundance barplots highlighting the taxa identified at different read depths(10k-100k reads). **(D)** Violin plots with overlaid boxplots showing the distribution of abundant and rare ASVs across subsampled read depths (10K-100K reads).

To directly assess the impact of sequencing depth on taxonomic recovery, datasets were subsampled across depths ranging from 10K to 100K reads/sample, followed by re-execution of the complete analytical workflow. Increasing sequencing depth resulted in a progressive increase in detected ASVs, rising from approximately ∼300 ASVs at 10K reads to ∼600 ASVs at 100K reads (Figure 4B). However, these gains were not accompanied by major restructuring of community composition. Across all evaluated depths, dominant gut-associated genera exhibited highly concordant abundance profiles with preservation of overall ecological structure (Figure 4C), indicating that additional sequencing depth primarily contributed incremental taxonomic recovery rather than alteration of core microbiota composition.

Analysis of rare taxa dynamics further demonstrated that increased sequencing depth predominantly expanded recovery of low-abundance community members (Figure 4D). Cumulative recovery of rare species increased progressively with sequencing depth, while their average relative abundance remained consistently low, indicating expansion of the rare biosphere rather than enrichment of dominant taxa. Per-sample analyses similarly demonstrated a depth-dependent increase in rare species detection across the cohort, with gains becoming progressively marginal at higher sequencing depths. Collectively, these findings indicate that approximately 40K-50K reads/samples capture the majority of biologically relevant diversity and species-level community structure, whereas sequencing beyond this threshold predominantly improves recovery of progressively rarer taxa without altering the composition of the abundant taxa.

## **3.** Discussion

The present study demonstrates the analytical and translational utility of Kinnex-enabled PacBio HiFi full-length 16SrRNA (V1–V9) sequencing for high-resolution microbiota profiling through a comprehensive evaluation spanning controlled microbial communities, technical replicates, and human fecal microbiota. While full-length sequencing has increasingly been recognized for its improved taxonomic resolving power, systematic assessments integrating analytical validation, reproducibility, real-world deployment, and implementation guidance remain limited. The current work addresses these dimensions within a single framework and provides evidence supporting the practical deployment of species level resolution for microbiota profiling.

Using defined positive-control communities, the workflow demonstrated accurate taxonomic recovery across varying community complexity and abundance gradients, including reliable discrimination of closely related taxa and reproducible detection of low-abundance members across dilution series. Importantly, these observations were reproduced across independent library preparations, sequencing pools, and SMRT runs, indicating stable performance throughout the workflow. Reproducibility across multiple technical layers is particularly relevant as microbiota studies increasingly expand toward large cohorts, longitudinal sampling, and translational applications where technical variance can substantially influence downstream interpretation.

Application with respect to healthy human fecal microbiota further highlighted the analytical value of full-length sequencing encompassing highly complex microbial ecosystems. The human gut microbiota is characterized by extensive phylogenetic density and intra-genus diversity, where conventional partial-region sequencing frequently limits interpretation at lower taxonomic ranks. In the present cohort, full-length sequencing enabled species-level characterization across dominant gut-associated genera and revealed substantial inter-individual heterogeneity while preserving overall ecological structure. In addition, recovery of multiple ASVs within prevalent taxa suggests that full-length profiling retains fine-scale sequence variation and captures additional layers of microbiota organization extending beyond conventional community-level summaries.

An important feature of this study was the internally controlled comparison between full-length V1–V9 and V3–V4 sequencing through computational reconstruction of V3–V4 datasets directly from identical full-length reads. This design isolates the effect of amplicon length independent of sequencing chemistry, sample processing, sequencing depth, and biological variability. Under these matched conditions, broad ecological patterns remained highly concordant between approaches, indicating that full-length sequencing preserves the biological signals captured by conventional workflows. However, substantial gains were observed at species level, particularly within taxonomically dense and clinically relevant genera including *Bifidobacterium, Prevotella, Blautia, Enterococcus,* and *Klebsiella*. These findings reinforce that the primary contribution of full-length sequencing lies not in altering community composition, but in extending interpretability through improved taxonomic resolution.

The study additionally provides practical guidance regarding sequencing depth requirements for 16SrRNA full-length microbiota studies. Integration of rarefaction analysis with explicit read-depth subsampling demonstrated that dominant community structure and most biologically relevant diversity were recovered at moderate sequencing depths, whereas deeper sequencing predominantly contributed to progressive recovery of low-abundance taxa and expansion of the rare biosphere. Importantly, the majority of core community structure and species-level signals were retained at approximately 20–30K reads per sample, while ∼50K reads per sample provided additional depth for robust species-resolved profiling and recovery of low-abundance bacteria. These observations have direct translational relevance, as they indicate that high-resolution full-length microbiota profiling can be achieved without ultra-deep sequencing. Under current high-throughput implementations and multiplexing configurations, such sequencing depths enable full-length profiling at per-sample costs approaching, and in some scenarios potentially comparable to, conventional short-read V3–V4 workflows while simultaneously delivering substantially greater taxonomic information content.

Collectively, the present work demonstrates that full-length 16S profiling can preserve the ecological structure captured by conventional short-read workflows while substantially expanding biological interpretability through species-level taxonomic resolution and recovery of ASV-scale diversity. By integrating controlled microbial communities, technical reproducibility analyses, real-world human gut microbiota profiling, comparative benchmarking against V3–V4 sequencing, and sequencing-depth optimization, this study provides a comprehensive framework for practical implementation of species-resolved microbiome investigations. As microbiome research increasingly progresses from descriptive community profiling toward mechanistic understanding of host–microbe interactions and clinically relevant microbiome phenotypes, the ability to accurately resolve closely related microbial taxa is expected to become increasingly important. Importantly, the combination of species-level resolution, moderate sequencing-depth requirements, and highly multiplexed full-length sequencing supports deployment across large population cohorts, longitudinal studies, microbiome biobanks, and public-health initiatives. Together, these findings position full-length 16S sequencing as an enabling technology for the next generation of microbiome research, where species-level microbial ecology can be interrogated at scales previously reserved for lower-resolution profiling approaches.

## CRediT authorship contribution statement

Conceptualisation was contributed by Devraj J. Parsanannanvar and Sridhar Sivasubbu. Methodology was contributed by Paras Sehgal and Ashriti Chettri. Investigation was carried out by Parth Sarin, Paras Sehgal, Sakshi Rai, Ashriti Chettri, and Rahul C. Bhoyar. Formal analysis was performed by Paras Sehgal and Vasudev Paveri. Data curation was undertaken by Parth Sarin and Paras Sehgal. Validation was performed by Parth Sarin, Sakshi Rai, and Ashriti Chettri. Visualisation was contributed by Vasudev Paveri. Resources were provided by Rahul C. Bhoyar, Shahzad Mirza, Rajesh Karkaryate, and Sridhar Sivasubbu. Writing of the original draft was undertaken by Parth Sarin and Paras Sehgal. Manuscript review and editing were performed by Parth Sarin, Sakshi Rai, Devraj J. Parsanannanvar, and Sridhar Sivasubbu. Supervision was provided by Devraj J. Parsanannanvar and Sridhar Sivasubbu. Project administration was carried out by Devraj J. Parsanannanvar. Funding acquisition was contributed by Sourav Sen Gupta and Devraj J. Parsanannanvar.

## Data availability

The data with respect to the paper will be made available to anyone with a reasonable request to the corresponding author of paper.

## Declaration of Generative AI and AI-assisted technologies in the writing process

During the preparation of this work, the author(s) used ChatGPT / Version 4.0] in order to Refine and improve the readability of the document. After using this tool/service, the author(s) reviewed and edited the content as needed and take(s) full responsibility for the content of the publication.

## Acknowledgements

The author thanks all the contributors of the study. Our work was supported by the Indian Council of Medical Research, the Department of Health Research, the Ministry of Health and Family Welfare. The authors would also like to acknowledge Mr. Harish Ramanathan for his assistance in establishing the computational infrastructure and bioinformatics environment required for data analysis. We also thank Dr. Bani Jolly and Dr. Vinod Scaria for their valuable technical discussions, scientific guidance, and constructive inputs during the design and interpretation of this study.

## Acknowledgement

The authors would like to acknowledge Mr. Harish Ramanathan for his assistance in establishing the computational infrastructure and bioinformatics environment required for data analysis. We also thank Dr. Bani Jolly and Dr. Vinod Scaria for their valuable technical discussions, scientific guidance, and constructive inputs during the design and interpretation of this study.

